# DeepMPS: Development and validation of a deep learning model for whole slide image base prognostic prediction of low grade Lung adenocarcinoma patients

**DOI:** 10.1101/2022.12.27.522072

**Authors:** Akshay Jagajampi, Seema Khadirnaikar, Prabhat S Malik, Deepali Jain, MB Naveen, Sudhanshu K Shukla

## Abstract

Lung adenocarcinoma (LUAD) is one of the most common cancers, and patients’ prognostication is crucial for treatment decisions. Histopathological images are the most generally accessible clinical information, however they have not been employed in clinical settings for prognosis. **In t**his study, we used WSIs and clinical data from TCGA (training and testing) and East Asian Cohort (EAS, Validation) to develop and validate DL-based prognosticator. To circumvent the need for manual ROI generation, WSIs from these patients were divided into smaller patches and scored. DeepMPS prediction model was built using the top scoring 226,383 patches. The DeepMPS model showed a C-index of 0.638 in the TCGA training cohort. The univariate and multivariate cox regression analysis identified DeepMPS as an independent predictor of survival (HR: 9.48, p-value: <0.0001) in the training cohort. The training cohort of patients was separated into low and high-risk groups at various points. Kaplan-Meier analysis showed the highest difference in survival of low and high-risk patients at the 75th percentile (HR: 3.58, 95% CI: 2.57-5.00, p-value: <0.0001). At the same cut-off as the training samples, TCGA testing cohort patients demonstrated a significant difference in survival when split into low and high-risk patients (HR: 2.30, 95% CI: 1.11-4.82, p-value: 0.044). The DeepMPS model was validated in the EAS cohort patients. DeepMPS risk score significantly segregated EAS patients into low and high-risk groups at the same cut-off point as the training cohort (HR: 2.09, 95% CI: 1.11-3.97, p-value: 0.008). In multivariate Cox regression analysis, the DeepMPS score outperformed the stage in survival prediction. We also compared the DeepMPS model with the previously developed DL-based model to show that it was the best predictor of survival with the highest C-index. In conclusion, we developed a robust DL-based prognostic model which can predict the LUAD outcome without manual intervention using histopathological images.

**Author Summary:** Right from the initial step of cancer diagnosis, histopathological images are the widely relied source of information for therapeutic decision making. Though these images carry a substantial amount of information, their use remains restricted to determining the grade of the tumor by pathologists. The advent of computational techniques has given rise to the ability to capture vital information besides the grade of the tumor, which humans might not be able to quantify visually. To this end, this work proposes a deep learning based model, Deep Multi-Modal Prognosis System (DeepMPS), to predict the prognosis of the patients based on the histopathological images to provide additional information to the pathologist which will aid in clinical decision making. DeepMPS predicts a risk score associated with each individual based on histopathological images and clinical factors. The risk score obtained from the proposed system is an independent predictor of survival. As the proposed system is independent of the manual region-of-interest (RoI) generation, it will ease the pathologist’s workload.

## Introduction

Lung cancer is the leading cause of cancer related deaths. In USA, lung cancer accounts for nearly 21% deaths per year which is more than combined deaths caused by colon, breast, and prostate cancers [1]. Lung adenocarcinoma (LUAD) is most common subtype of the lung cancer with ever increasing incident. The diagnosis of the disease is done based on histopathological image analysis and clinical staging system has been standard for determining the prognosis. Histopathological features further refine the prognostication and therapeutic decision making [2]. Current standard for early stage LUAD is surgical resection with or without adjuvant systemic treatment [3]. The adjuvant treatment decision is largely based on clinical and pathological criteria. Further attempts of refining these criteria using genomic strategy has been successful but wider clinical application is limited due to technical requirements and cost [4].

Whole slide images (WSI) are histopathological images captured using high end microscopic method. As WSI contains images data of full slide it contains more information about the disease apart from pathological subtypes of cancer. These histopathological images are used for the classification of LUAD patients into different subtypes, but can also be used for prognostic stratification and therapeutic decision making [5]. Hence, any prognostic model developed using histopathological images are easily adoptable in clinical practice. Despite its clinical utility, WSIs are not used for the prognostic determination of the LUAD patients. This is due to impossibility of identifying minor details of the image with human eyes. To this end, deep learning (DL) methods have shown us tremendous improvements in identification, classification, segmentation and survival analysis of various malignancies [6, 7].

During the last few years, many DL-based methods have been proposed for survival prediction using various data modalities, including gene expression, methylation, radiomics images, and WSIs. Recently, research suggested the Automatic Cancer Detection and Classification in whole-slide Lung Histopathology challenge to evaluate several computer-aided diagnoses (CADs) approaches to the automatic detection of lung cancer [8]. Another study has used Convolution neural network (CNN) to identify the cell types in LUAD patients [9]. In colorectal cancer, DL methods have been used to predict the status of molecular pathways and mutations using WSIs [10]. Another study has developed DL based method for the survival prediction of colorectal cancer [11]. A CNN-based method has been developed to identify the early stage LUAD, which also predicts the prognosis indirectly [12]. However, to the best of our knowledge, there are no studies proposed, developed, and validated in the direction of direct LUAD survival prediction. Here, we have used two independent datasets to develop and validate DL based method for LUAD patients’ survival prediction. We have also shown that our model outperforms the previous models and clinical factors like stage and age are significant for the survival prediction. The proposed model is validated in an independent external validation dataset as well.

## Material and Method

### Data Cohorts

We have used two independent cohorts of patients for this study. The first cohort corresponded to the TCGA dataset consisting of 330 early-stage LUAD data (stages I, II, and III), which was used for training, validation and testing our DL model. The dataset included each patient’s WSIs and other clinical data (age, stage, gender, smoking habit, etc.). Information from WSIs and clinical data was used as multi-modal inputs to train the DL model. The second cohort corresponded to the East Asian Sample (EAS) cohort, which consisted of WSIs and clinical data for 190 LUAD patients [13]. This was used for independent validation, i.e., our DL model had not encountered this dataset during its training, validation, or testing. The details of these two datasets are summarized in Table 1.

**Table 1:** Summarizing the characteristics of patients in TCGA-LUAD and EAS cohorts.

| Factor | TCGA Training Cohort | TCGA Validation Cohort | EAS validation cohort | P-value |
| --- | --- | --- | --- | --- |
| No. of patients | 330 | 86 | 190 |  |
| Age, y, mean (SD) | 66.3 (9.44) | 64.2 (11.44) | 49.63 (29.68) | 0.07<sup>#</sup> |
| Female sex, No. (%) | 182 (55.1) | 46 (53.48) | 105 (55.2) |  |
| Median survivor follow-up, mo. | 22.17 | 22.13 | 41.98 |  |
| Median Survival, months | 50.03 | 49.73 | Undefined |  |
| Censored samples | 219 | 57 | 141 |  |
| Samples with Event | 119 | 29 | 49 |  |
| Stage No. (%) |  |  |  |  |
| I | 192 (58.2) | 52 (60.5) | 117 (51.6) | 0.07<sup>*</sup> |
| II | 87 (26.3) | 24 (27.9) | 36 (18.9) |  |
| III | 51 (15.5) | 10 (11.6) | 37 (19.4) |  |

### Image pre-processing

One of the major challenges in using data from WSIs in our DL model is that they are large (on an average 1 GB) and they vary in size (10 MB to 3.4 GB). The utilization of these WSIs in their entirety is impossible due to the high cost of training a DL model, and the background of WSIs does not contain any useful information. Hence, we employed various image pre-processing techniques in order to make the images usable for our analysis (**Figure 1a)**. Firstly, WSIs were scaled down by a factor of 32 and used as a reference for preprocessing tasks, including eliminating the background, creating patches, and scoring the patches. Since the background of WSI was mostly white, we generated masks to eliminate this region and obtain the ROI (method detailed in Supplementary Methods: Mask generation). The masked image thus generated was further used to generate the tissue patches with the background ignored (Supplementary Methods: Generation of Tissue Patches). Next, to further filter the number of patches, we assigned scores to patches based on tissue percentage and color characteristics as explained in Supplementary Methods: Selection of Tissue Patches. A WSI with selected ROI patches is shown in Supplementary Figure S1. The top 650 patches were then selected for further pre-processing, with blurred patches and ink-mark patches removed. A total of approximately 226,383 patches spanning all the WSIs were selected based on the proposed pre-processing technique, out of which 70% of patches were used for training, 15% of patches for validation, and the rest 15% for testing. Finally, all the patches were resized to 300 × 300 to be compatible with the standard input size for the pre-trained model we used in our study [14]. Moreover, we used clinical data such as age, gender, and cancer stage in addition to the image data from WSIs [15]. Gender, and cancer stage being categorical data, were converted to numerical values using one hot encoding technique.

**Figure 1:**
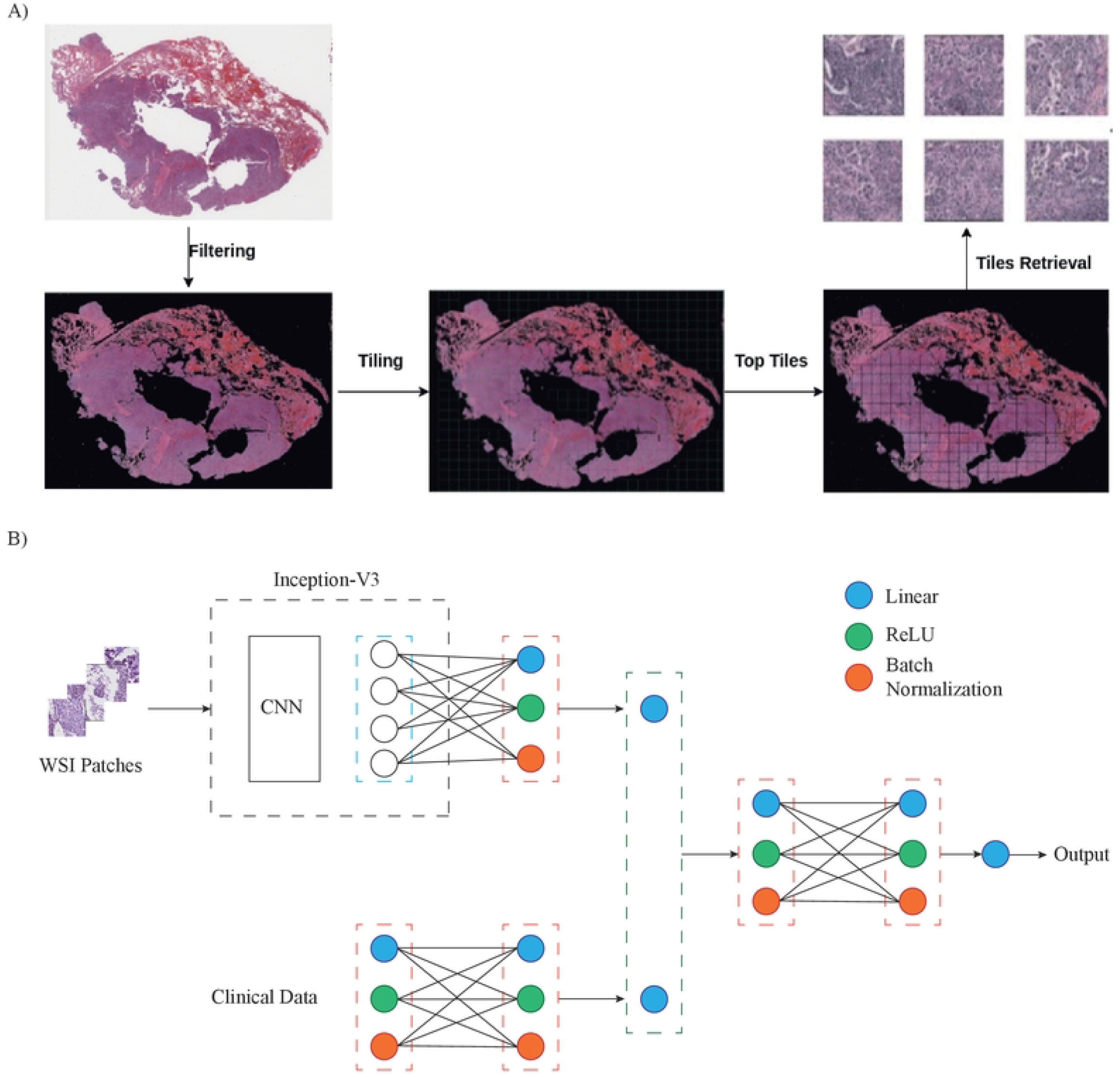
a) Overview of pre-processing tlie whole slide image (WSI). b) Overview of the proposed DEEPMPS that mainly consisted of IV3 and multiple FFNNs

### DL Model

In this work, we introduce a Deep Multi-Modal Prognosis System (DeepMPS) that consists of an existing pre-trained Inception-V3 (IV3) model (a proven network for image classification) and a fully connected feed-forward neural network (FFNN). The multimodal input design pattern addresses the problem of representing different data types (in this case, image and clinical data) that can be expressed in complex ways by concatenating all the available data representations.

Initially, a pre-trained IV3 model was used, and the final layers were fine-tuned using the selected patches as complete training of the model on image data was time-consuming and resource intensive (Figure 1 b). However, the performance of this model was not satisfactory (Figure 1 c and d). Hence, the IV3 model was trained from scratch. Also, the final layers of IV3 were modified to reduce the number of parameters, thereby reducing the training time. This further allowed the model to learn more appropriate features with respect to LUAD images during training. We added two linear layers where the first layer consisted of 512 neurons and the final output layer consisted of 8 neurons. Similarly, a second FFNN consisted of three sets of linear layers followed by a ReLU activation layer and batch normalization layer. The final layer of the second network also consisted of 8 neurons. We concatenated the output feature vectors of these two networks to feed it as an input to the final fully connected FFNN consisting of two sets of linear, dropout, and batch normalization layers (Figure 1 b).

Histopathology patches selected based on the score obtained by Supplementary Equation (S5) were fed as input to our modified IV3 model; additionally, the clinical data consisting of age, gender, and stage information were used as input to the second FFNN. Gender, and cancer stage being categorical data, were converted to numerical values using one hot encoding technique. Further, we modeled survival prediction as a ranking problem and the final linear layer would output a single value, which was treated as the risk score associated with the corresponding patient.

### Training

The TCGA-LUAD cohort was used to train the model. However, it was imbalanced and consisted of more patients belonging to the alive category than the dead category. As a result, data augmentation techniques such as flip, rotation, and sharpening were used to obtain a balanced dataset. As the data available to train the model in terms of the number of patients was less, there was a high probability of the model overfitting the data. Hence, the technique of K-fold cross-validation was used during training. We first split the data into training, validation, and testing to implement this method in a 70:15:15 ratio. The variance of the resulting estimate reduces as K increases but training such an algorithm is computationally intensive as it should be trained K times from scratch. To balance these two factors, we chose K = 6 for our K-fold cross-validation. In this method, one of the six-folds is used for validation, and the remaining five-folds were used for training where every data point gets to be used for training 5 (i.e., K-1) times and exactly once for validation. This helped to provide an unbiased evaluation of the model’s fitness.

At the final layer, the predicted risk score had features coming from both image and clinical data networks. Note that the weights were updated in all three networks during back-propagation, with weights being updated independently in the first two networks (Figure 1 b). We used a loss function based on the Cox partial likelihood (Cox-PH). Cox-PH model accounts for the censored data and simultaneously models the effect of multiple factors on survival (method detailed in Supplementary Methods: Loss Function). The steps involved in tuning the hyper-parameters used to train the DL model are summarized in Supplementary Methods: Tuning Hyperparameters. Training and validation of the DL model based on a six-fold crossvalidation method took approximately 170 hours on NVIDIA V100 single GPU.

We use the Concordant index (C-index) our metric for performance evaluation (Supplementary Methods: C-index). It is the most widely adopted and a standard evaluation metric to evaluate the performance of survival prognosis models [11, 15].

### Kaplan-Meier and Cox regression analysis

Kaplan-Meier analysis was used to show the difference in survival of different groups. The significance of survival difference was tested using the Log-rank test in *GraphPad version 9.2* software (San Diego, CA). The association of various factors with survival was performed using univariate and multivariate Cox regression analysis in *Survival* package of R.

## Results

### Patients Characteristics

We utilised 330 patients data from the TCGA-LUAD cohort of samples for training and internal validation and 190 patients data from EAS cohort for external validation. The clinical characteristic of these patients are given in table 1. There was no statistically significant difference in clinical characteristic of patients of these cohorts.

### Development of DL based prognostic model

Initially we used our network (Figure 1 b) with the pre-trained IV3 model and checked the WSI patch regions it focused on using the Grad-CAM [16]. We discovered that the derived features of WSIs using a pre-trained model did not accurately represent the patch information throughout our experiment (Figure 1 c). As a result, we trained our model from scratch for both L1 and L2 regularization, which not only improved the accuracy but also its performance, as demonstrated in Figure 1 d.

The L2 regularization was used in our final model since the DeepMPS with L2 regularization performed better than the DeepMPS with L1 regularization. Since we used sixfold cross validation, each fold gets a chance to appear in the training set five times, ensuring that every observation in the dataset appears in the training dataset, allowing the model to learn the underlying data distribution better. As expected, the loss decreases and C-Index increases for all the six folds throughout training and validation over 120 epochs (Supplementary Figure S3 A, B, and C). Additionally, we discovered that training a pre-trained IV-3 model with the final few modified layers resulted in a significantly larger training loss than training a model from scratch, (**Supplementary Figure S3 A and D**). This was also confirmed by the settling of C-index values, which are significantly lower while training the final few layers of a pretrained model than when training a model from scratch (**Supplementary Figure S3 B and E**). In the end, the trained model weights for each fold were used to evaluate the model’s performance on unseen TCGA test data and an independent EAS dataset. The model worked well in both cohorts, indicating that training the model from scratch using both image and clinical data generalizes well in both the cohorts (**Table 2).**

**Table 2:** The proposed model’s C-index values for all six folds for both the TCGA and EAS lung cancer datasets.

| $k^{\text{th}}$-Fold | TCGA - LUAD (median) | EAS |
| --- | --- | --- |
| 1 | 0.609 | 0.557 |
| 2 | 0.678 | 0.566 |
| 3 | 0.586 | 0.558 |
| 4 | **0.638** | **0.648** |
| 5 | 0.635 | 0.502 |
| 6 | 0.664 | 0.517 |

### Risk Score generation

The DeepMPS produced a continuous output which was treated as a risk score associated with the patient, with a high-risk score indicating a lower chance of survival and a low-risk score indicating a higher chance of survival. To understand the prognostic value and independence of the DeepMPS risk score over other clinical parameters, we performed univariate and multivariate Cox regression analysis with stage and DeepMPS risk score as co-factor in both TCGA training and testing separately, and as well as the combined data. This analysis showed that risk score is independent and strongest predictor of survival in TCGA dataset (**Table 3**). **To categorise patients into low and high risk groups, we** dichotomized DeepMPS risk score at various cut-offs and identified 75^th^ percentile as optimum in training cohort (**Figure 2 A**). In TCGA training samples, patients with normalised risk score of more than 0.54 had statistically significantly worse overall survival (**HR: 11.28, 95% CI-4.32 to 29.42, p-value:<0.0001**, **Figure 2 B**). Similar to training samples, patients with >0.54 normalised risk score in TCGA validation samples also showed significantly poor survival when compared to patients with low risk score (**HR: 2.31, 95% CI-1.11 to 4.82, p-value:0.044, Figure 2 C and D**). Furthermore, the risk score was able to divide the full TCGA cohort samples into low and high risk with statistically significant difference in survival (**HR: 3.58, 95% CI-2.57 to 5.00, p-value:<0.0001, Figure 2 E and F).**

**Table 3:**
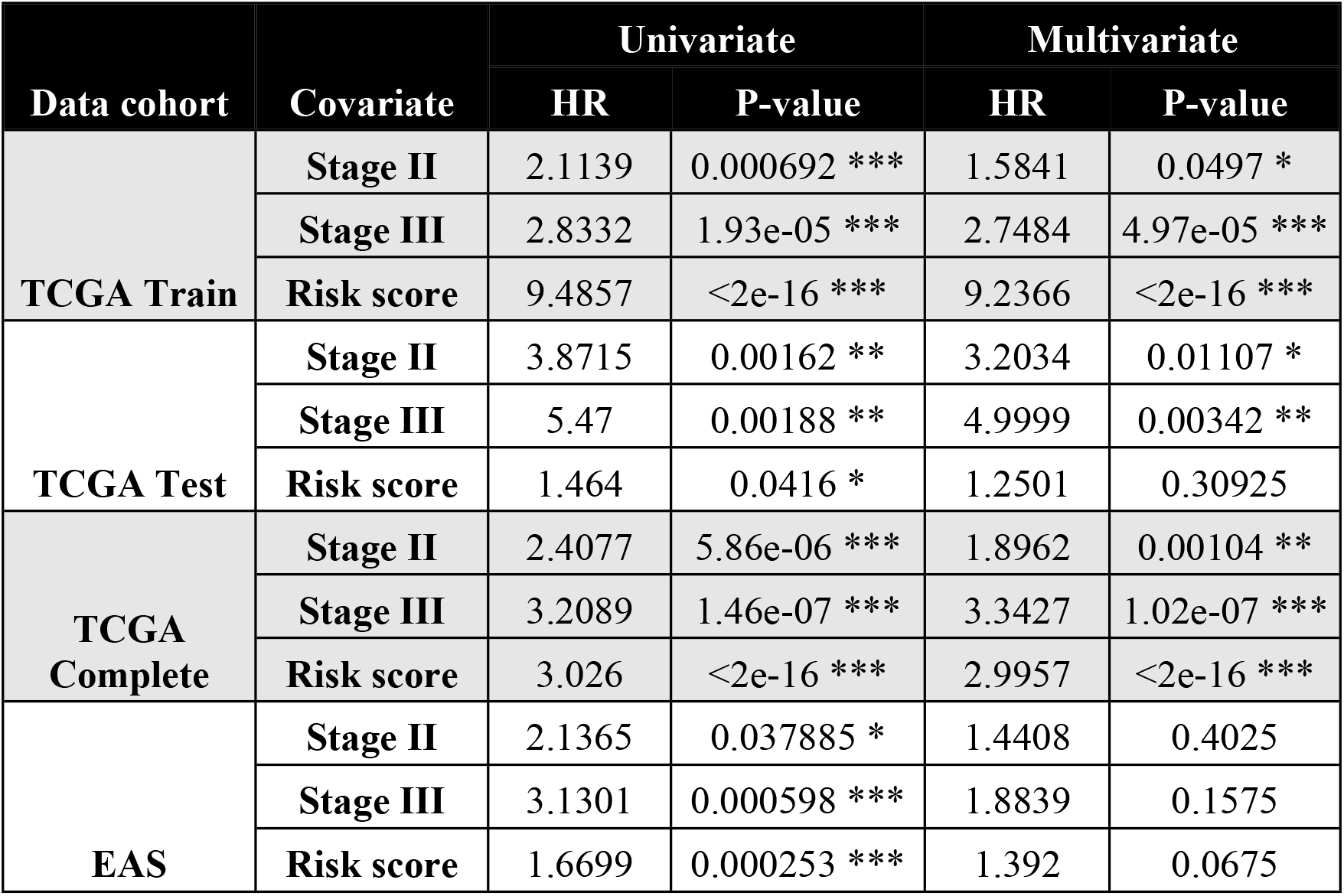
Summarizing the univariate and multivariate Cox regression results in TCGA and EAS cohorts.

| Data cohort | Covariate | Univariate |  | Multivariate |  |
| --- | --- | --- | --- | --- | --- |
|  |  | HR | P-value | HR | P-value |
| TCGA Train | Stage II | 2.1139 | 0.000692 *** | 1.5841 | 0.0497 * |
|  | Stage III | 2.8332 | 1.93e-05 *** | 2.7484 | 4.97e-05 *** |
|  | Risk score | 9.4857 | <2e-16 *** | 9.2366 | <2e-16 *** |
| TCGA Test | Stage II | 3.8715 | 0.00162 ** | 3.2034 | 0.01107 * |
|  | Stage III | 5.47 | 0.00188 ** | 4.9999 | 0.00342 ** |
|  | Risk score | 1.464 | 0.0416 * | 1.2501 | 0.30925 |
| TCGA Complete | Stage II | 2.4077 | 5.86e-06 *** | 1.8962 | 0.00104 ** |
|  | Stage III | 3.2089 | 1.46e-07 *** | 3.3427 | 1.02e-07 *** |
|  | Risk score | 3.026 | <2e-16 *** | 2.9957 | <2e-16 *** |
| EAS | Stage II | 2.1365 | 0.037885 * | 1.4408 | 0.4025 |
|  | Stage III | 3.1301 | 0.000598 *** | 1.8839 | 0.1575 |
|  | Risk score | 1.6699 | 0.000253 *** | 1.392 | 0.0675 |

**Figure 2.**
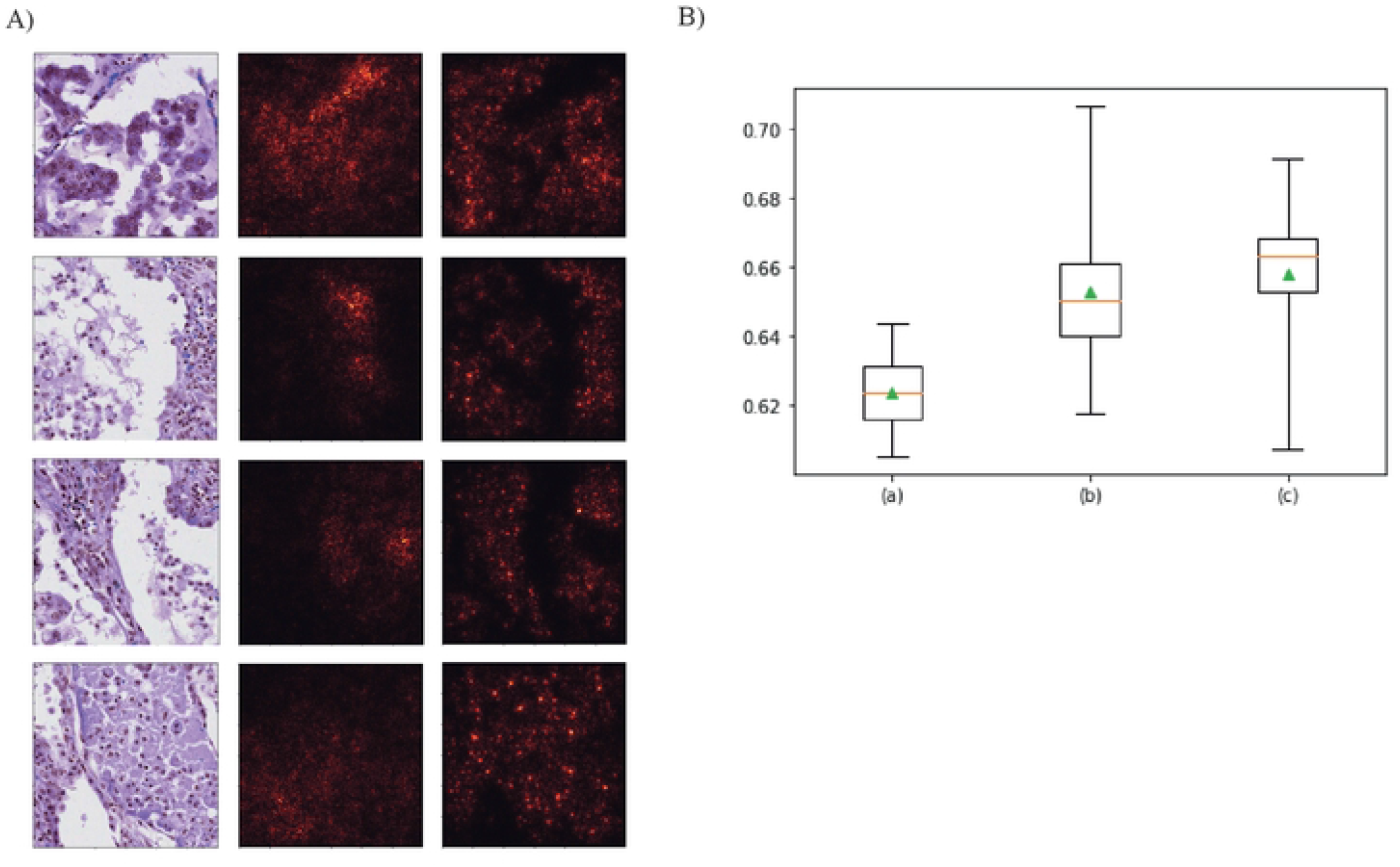
A) Original tissue patch images (leftmost) along with the corresponding Grad-CAM (GCAM) hcatmap images of a pre-trained model (middle) and a model trained from scratch (rightmost). B) Box-plots showing C-index of (a) Pre-trained model with median C-index = 0.625. (b) The model trained from scratch with L1 regularization and a median C-index = 0.65. (c)The model trained from scratch with L2 regularization and a median C-index = 0.667.

**Figure 3:**
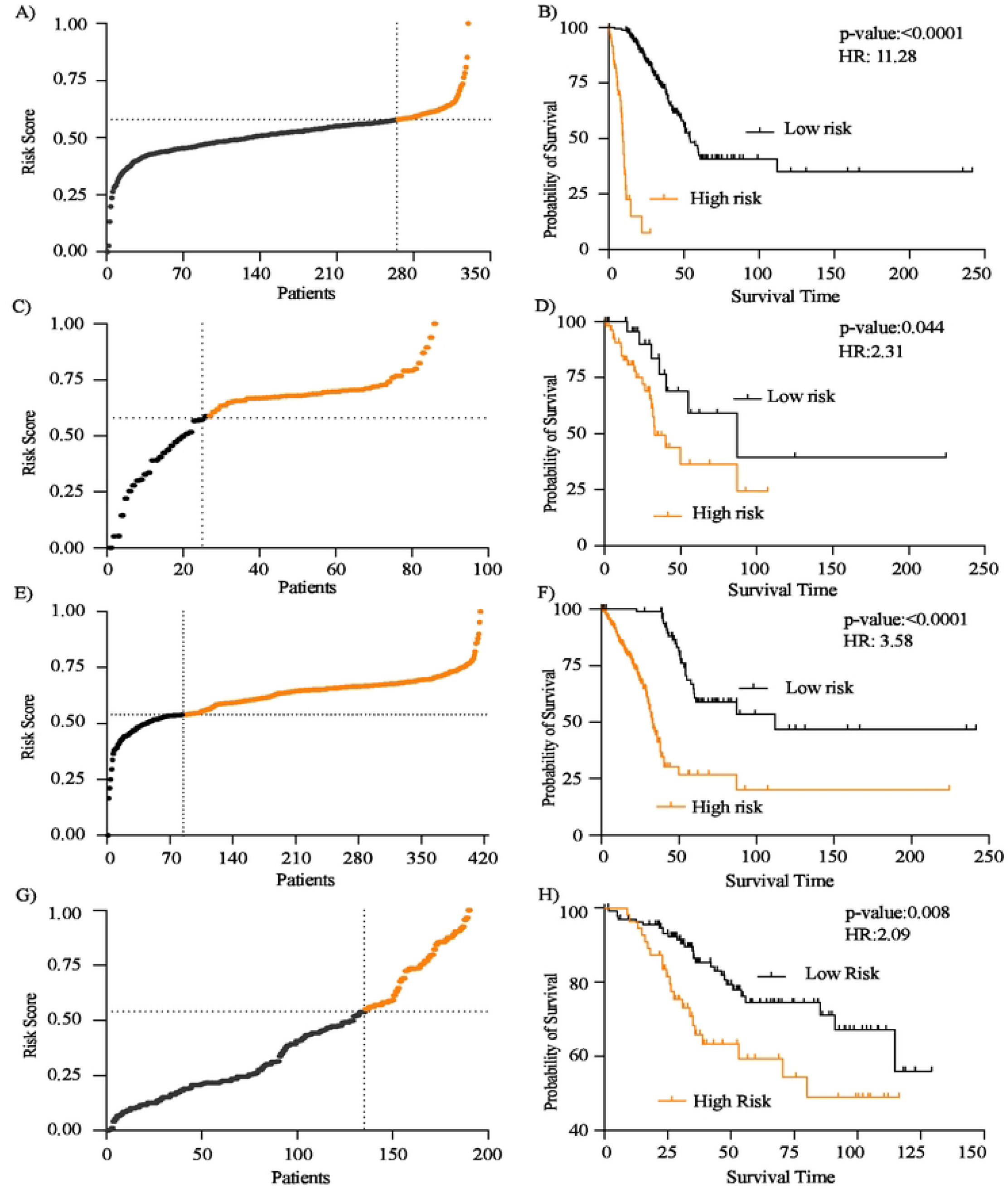
Risk score and KM survival curves A) The risk score distribution associated with patients in the TCGA train cohort. The dashed line represents the 75th percentile, which was used to categorize patients into high and low-risk groups. B) The KM plot shows the high-risk groups (orange), and low-risk groups (black) with p ≤ 0.05 indicating that the two groups differ significantly in TCGA train dataset. C) The risk score distribution associated with patients in the TCGA test cohort. The dashed line represents the 75th percentile in the training dataset, which was used to categorize patients into high and low-risk groups. D) The KM plot shows the high-risk groups (orange), and low-risk groups (black) with p ≤ 0.05 indicating the significant difference in survival in both groups in TCGA test dataset. E) The risk score distribution associated with patients in the complete TCGA cohort. The dashed line represents the 75th percentile in the training dataset, which was used to categorize patients into high and low-risk groups. F) The KM plot shows the high-risk groups (orange), and low-risk groups (black) with p ≤ 0.05 indicating the significant difference in survival in both groups. G) The risk score distribution is associated with patients in an independent EAS cohort. The horizontal dashed line is the threshold (taken from the training set) value for categorizing patients into low and high-risk groups. H) A lower p-valuc (p ≤ 0.05) shows that the chosen threshold value was able to dichotomize patients into risk groups that arc significantly different in EAS cohort.

### Comparison of results

We compared the performance of DeepMPS model to that of existing machine learning based models using C-index values. **Table 4** shows that our proposed DeepMPS model outperformed all the other models. We also trained separate models on clinical and image data as part of the our analysis. We observed that the model trained solely on image data provides more insights into the data and produces better results (C-index = 0.62) than the model trained solely on clinical data (C-index = 0.47) (**Supplementary Table S1**). However, our proposed DEEPMPS model trained using both clinical and image data outperforms models trained solely on image or clinical data (C-index = 0.64 vs 0.62 and 0.47).

### Independent validation of risk score

Next, we used EAS cohort of 190 samples for independent validation of DeepMPS risk score as prognosticator (**Table 1**). The DeepMPS risk score was calculated using the same method as that of the training cohort. The distribution of risk score and relation with patients’ status are shown in **Figure 2 e and f**. Using the same cut off used in training samples, DeepMPS risk score was able to stratify the EAS cohort patients significantly (**HR: 2.31, 95% CI-1.11 to 4.82, p-value:0.044**) (**Figure 2 f**). These results show the robustness of the risk score as prognosticator. We also performed univariate Cox regression analysis and showed that risk score is strongest predictor of survival (**Table 3**). In multivariate analysis, stage was not found to be a predictor of survival (**Table 3**). In contrast, risk score was still better prognosticator with p-value approaching to significance (**Table 3**).

## Discussion

LUAD is a major cause of cancer mortality and morbidity throughout the world causing more than 150000 deaths in united states alone. There is a need for improvisations in the current clinical and pathological prognostic strategies for better therapeutic decision making. Although some clinically approved prognostic gene panels are available, the use is limited [17]. This suggest for further development of the prognostication strategies for LUAD patients. Using data from TCGA WSIs, we developed a deep learning based model for the prognosis prediction of LUAD patients. Our method is completely automated and does not require manual ROI generation by trained pathologist. We independently validated the prognostic model on a completely unseen cohort of 190 patients. Crucially, our model remain an independent predictor of overall survival in LUAD patients with more than two fold increased risk of death in high risk patients compared to low risk patients. Our model also utilises the other clinical data like stage and age for the improved prediction.

To the best of our knowledge, ours is the only study where WSIs are utilised without manual ROI generation. The DeepMPS score also showed high c-index and outermore the previously reported deep learning based models. Additionally, our model performed well irrespective of the WSI obtained from different geographical area, equipment’s used for scanning, and treatment methods.

Histopathological images including WSIs which are used for the diagnosis of the disease, are one of the most commonly and easily available clinical data. Hence, any prognostication model developed using histopathological images would be available for prognosis prediction of every patients without additional testing requirement. Recently, deep learning models generated using already available image data have developed a lot of interest. Computer algorithms have been in use since 1993 for image analysis for cancer grading, diagnosis and prognostic predictions [18]. Recently, deep learning based algorithms have dramatically improved the results of image analysis of pathological images [19–21]. Many DL based model have been developed to use for histopathological image analysis in lung cancer. A CNN based model was developed to differentiate between cancerous and non-cancerous tissue with an accuracy of 89.8% in testing set [22]. This study also proposed the prognostic prediction of patients using tumor shape and boundary features. However, this study required selection of malignant and non-malignant region identification by pathologist. Similarly, Coudray et al have used deep learning based method to automatically classify LUAD, LUSC and normal tissue [23]. Further, authors also identified mutations like STK11, TP53, EGFR etc. in non-small cell lung cancer using histopathological images. Peter et al used DL based method to develop QuPath to analyse the histopathological images [24]. In 2019, Wang et al developed ConvPath based on convlolution neural network to identify various cell types like tumor cell, stromal cell, and lymphocyte [9]. Authors have used the cell type information to developed prognostic model for prognostication of NSCLC patients. However, published models have inherent shortcomings, including requirement for ROI generation, non-validation on external dataset. Our method overcomes these limitation and works on external unseen data with no manual ROI requirement which makes it’s wider clinical application possible.

One of the shortcomings of our research is that it does not include another sample cohort for the validation of the model. In addition, we believe that if a greater number of examples were used to train our model, it would be able to perform even better.

Taken together, the novelty of DeepMPS lies in our approach to automatically select the ROI. Also, we have used scoring method to use patches of WSIs with high tissue and tumor content in an unbiased manner. The other crucial part of our study is validation of model in an independent dataset generated using DeepMPS of different ethnicity treated with different treatment protocol. We believe that DeepMPS has potential to automate the prognostic prediction of LUAD patients with minimal computational power.

## Authors Disclosers

None to disclose.

## Authors Contribution

Study Conception and design: AJ, NMB, SKS

Data collection: AJ, SK, NMB

Analysis and interpretation of results: AJ, SK, NMB, SKS

Draft Manuscript Preparation and editing: AJ, SK, PSM, DJ, NMB, SKS

## Acknowledgement

The results described in this publication were drawn in whole or in part from data generated by the TCGA research network. We would like to express our appreciation to AnantGanak, the High Performance Computing facility at IIT Dharwad.

